# Biological and statistical interpretation of size-at-age, mixed-effects models of growth

**DOI:** 10.1101/845222

**Authors:** Simone Vincenzi, Dusan Jesensek, Alain J Crivelli

**Author notes:** **Authors’ Contributions** SV conceived the ideas, designed the methods, and run the analyses; AJC conceived and run the Marble trout Project and AJC and DJ collected and curated the data; SV analyzed the data; SV led the writing of the manuscript. All authors gave final approval for publication.

## Abstract

The differences in life-history traits and processes between organisms living in the same or different populations contribute to determine their ecological and evolutionary dynamics. Recent advances in statistical and computational methods make it easier to investigate individual and group variation in life-history traits.

We developed mixed-effect model formulations of the popular size-at-age von Bertalanffy and Gompertz growth functions to estimate individual and group variation in body growth, using as a model system four freshwater fish populations living in Slovenian streams, where tagged individuals were sampled for more than 10 years. We used the software Template Model Builder to estimate the parameters of the mixed-effect growth models.

Estimates of asymptotic size from the Gompertz and von Bertalanffy models were not significantly correlated, but their predictions of size-at-age of individuals were strongly correlated (*r* > 0.99). Tests on data that were not used to estimate model parameters showed that predictions of individual growth trajectories using the random-effects model were accurately predicted (*R*^2^ > 0.80 for the best models over more than 500 predictions) starting from one single observation of body size early in life. Model results pointed to size ranks that are largely maintained throughout the lifetime of individuals in all populations.

## 1 Introduction

Understanding the causes of within- and among-population variation in vital rates of organisms, such as their probability of survival, growth, migration, and reproduction, life histories (i.e., how vital rates vary together and the tradeoffs among them), and population dynamics (i.e., how the number of individuals in a population changes over time) is a central topic in ecology and evolutionary biology.

Among vital rates, body growth is likely the one that has historically received the most attention, since survival, sexual maturity, reproductive success, and movement and migration are frequently associated with, or affected by, growth and body size [1]. Therefore, variation in growth within and among populations can greatly affect their ecological and evolutionary dynamics [2–4], and a better understanding of growth will always be an important problem in biology.

Mathematical and statistical models have been a staple of research in ecology and evolutionary biology for many decades; they have been used, among other goals, for statistical inference, identification, quantification, and prediction of biological and environmental processes and their realizations, and decision-making in species management and conservation [5]. When studying the growth of organisms in either applied or theoretical contexts, the identification of a functional form that can reasonably approximate the observed trajectories is a crucial first step in the development and application of a useful growth function. Structured or parametric models for growth of vertebrates imply a basic functional form: size increases monotonically with time, growth is faster early in life, and size usually tends to an upper asymptote later on in life. Since the modeling of mechanisms is more informative than curve fitting (e.g., it allows to predict outside of the range of data observed and test hypotheses on the determinants of growth), the parameters of the growth function should provide some physiological or life-history insights into the growth process. However, depending on their assumptions and parameterizations, growth functions fitted on the same data can provide different — sometimes in disagreement — insights into the growth process.

For vertebrates, the most widely used growth functions for size-at-age (the other macro-group of growth functions is auto-regressive functions, which use the individual’s previous size to predict its size at the next time point, see [6] for a modern study of those functions) have been the Gompertz [7, 8] and the von Bertalanffy growth functions [9], two special cases of the Richards growth function [10]. Both are sigmoid functions, a reasonable choice when modeling the growth of organism with slow initial growth, which then increases in speed before leveling off toward adult value. Their most popular parameterizations include three parameters, one of which — the asymptotic size — is common between the Gompertz and the von Bertalanffy growth functions.

In the vast majority of applications of growth models in ecology and evolutionary biology, the parameters of growth functions have been estimated at the population level, and interpreted as those of an average individual in the population [11]. However, estimates of parameters at the population level neither describe nor explain the large variation in growth often observed in organisms living in the same population; this greatly limits the scope and breadth of applications of the “average” growth functions in ecology and evolutionary biology. There are many examples (e.g., fish) of individuals living in the same population and of similar age with very different — sometimes hugely different — sizes [12, 13]; for most investigations, we are thus interested in each individual, and not population- or group-average, growth trajectory. Understanding which biological and environmental processes determine group and individual variation in growth may also improve predictions of future growth and size of individuals and populations, which is valuable for the conservation and management of species [14].

However, understanding the nature and contribution of sources of variation in growth faces a number of experimental, methodological, and computational challenges. First, we need longitudinal data (i.e., multiple observations of the same individual through time) to tease apart growth variation emerging from persistent differences among individuals or among groups from variation due to stochastic processes [6, 15], and to predict individual and average group growth trajectories. Especially for long-lived and elusive organisms such as fish, the collection of longitudinal data can take many years and much effort (individuals are usually tagged with a unique ID identifier), when at all possible. As marine scientist John Sheperd said: “Managing fisheries is hard: it’s like managing a forest, in which the trees are invisible and keep moving around” (http://jgshepherd.com/thoughts/).

Longitudinal data are thus often sparse, and data for a particular individual are unlikely to be sufficient for the estimation of parameters of the growth model for that individual. However, we may leverage data of other individuals that are thought to be similar to support the estimation of model parameters for the data-poor individual (i.e., the concept of “borrowing strength”, [16]). Models in which all members in a group influence the estimate of each effect are called either hierarchical, random-effects, multi-level, or mixed-effects models [17].

Random effects in mixed-effects models are realizations of a stochastic process. The assumption of common statistical distribution implies a dependence between random effects, which means that the estimate of the random effect for an individual is influenced by data and estimates of random effects for all other individuals relating to the same factor or group (say, for an organism, year-of-birth, sex, location, or, at a coarser grain, the whole population). Since in mixed-effects models those realizations will be pulled toward the mean of the group (“shrinkage”, [16]), the realizations that are strongly supported by data contribute more information to the statistical distribution of the effects, and we are then less likely to over-interpret processes that are the result of small sample sizes. Modeling and estimating random effects also address the lack of independence between repeated measurements of the same individuals [17]. Mixed-effects models of growth thus provide an intuitive framework for estimating heterogeneity of growth among individuals or groups of individuals, approximate their growth trajectories, and predict their future sizes [18].

Second, growth models of organisms are often — if not always — non-linear, and the estimation of parameters of non-linear models is intrinsically more computationally demanding than for linear models. However, fast and reliable approaches to parameter estimation are needed to investigate multiple parameterizations of complex models [11]. Although parameter estimation in mixed-effects models can still require considerable work, the many — and in some cases groundbreaking — advances in theory, algorithms, and software of the last few years have made the development of mixed-effect models and the estimation of their parameters much easier (i.e., the optimization algorithms more reliably converge) and faster than it used to be [19, 20].

Here, we take advantage of those recent theoretical and computational advances to propose mixed-effect formulations of the von Bertalanffy and Gompertz growth functions that can help model individual and group variation in the growth of organisms. Our work on growth is motivated by the long-term study of populations of marble trout (*Salmo marmoratus*), brown trout (*Salmo trutta* L.), and rainbow trout (*Oncorhynchus mykiss*) living in Slovenian streams. Marble, brown, and rainbow trout populations were sampled each June and September for more than ten years (2004-2015), and the data show substantial variation in size-at-age of individuals [21–23]. [24] and [23] showed that fast-growth allows trout populations to recover after massive mortality events, such as those caused by flash floods, since fecundity tends to increase with fish size. In addition, since growth and body size are heritable [25], there is potential for the evolution of growth rates toward faster growth in a population affected by massive mortality events [26, 27]. Therefore, models of growth that can predict growth trajectories for individuals, groups (e.g., born before or after extreme events), and populations will greatly benefit our understanding of the resilience of populations to extreme events and of the evolution of body growth in the affected populations.

In this paper we: (1) propose mixed-effects, individual-based formulations of the well-known and widely used von Bertalanffy and the Gompertz growth functions, and show how their parameters can be efficiently estimated with Template Model Builder, an open-source statistical software package for fitting non-linear, mixed-effects statistical models using the Laplace approximation [28]; (2) use four salmonid populations living in Slovenian streams as a model system to test the ability of the mixed-effects growth models (i.e., von Bertalanffy and Gompertz growth functions with predictors of their parameters, including random effects) to predict unobserved size-at-age of individuals; (3) test whether the life-history and biological processes determining the observed individual variation in growth (in particular the maintenance of size hierarchies among individuals throughout their lifetime), are inferred differently when using the von Bertalanffy or the Gompertz growth function.

## 2 Material and Methods

### 2.1 Study area and species description

Our system comprises the marble trout (*Salmo marmoratus*) populations of Lower Idrijca [LIdri_MT] and Upper Idrijca [UIdri_MT] [21], rainbow trout (*Oncorhynchus mykiss*) population of Lower Idrijca [LIdri_RT] [29], and brown trout (*Salmo trutta* L.) population of Upper Volaja [UVol_BT] [22]. All streams are located in Western Slovenia [29]. The streams Lower Idrijca and Upper Idrijca are a few hundred meters apart and are separated by a dam. In LIdri, marble trout [LIdri_MT] live in sympatry with rainbow trout [LIdri_RT] [29, 30]. Both UIdri_MT and UVol_BT live in allopatry, that is no other fish species live in the streams. The LIdri_RT population was created in the 1960s [29] and the UVol_BT population in the 1920s [22] by stocking rainbow and brown trout, respectively. Both populations have been self-sustaining since their inception. Further details on the demographic and life-history traits of these salmonid populations, along with their conservation status and current management practices, can be found in [29].

### 2.2 Sampling

Populations were sampled bi-annually in June and September. The first sampling for LIdri_MT, LIdri_RT, and UIdri_MT was in June 2004, and in September 2004 for UVol_BT. If captured fish had length *L* > 115 mm, and had not been previously tagged or had lost a previously applied tag, they received a Carlin tag [31], and their age was determined by reading scales. Genetic tagging was then used to identify fish that had received a new tag after losing theirs between sampling occasions [29]. Fish are 0+ (juveniles) in the first calendar year of life, 1+ in the second year and so on. Sub-yearling marble, rainbow, and brown trout are smaller than 115 mm in June and September, so fish were tagged when at least aged 1+. Sampling protocols are described in greater details in [21] and [22]. Total number of fish aged 1+ or older sampled from 2004 to 2015 was 1371 for LIdri_MT, 670 for UIdri_MT, 250 for LIdri_RT, and 2636 for UVol_BT.

### 2.3 Growth functions

Here, we introduce two of the most widely used growth functions in ecology, evolutionary biology, and fishery science: the von Bertalanffy and the Gompertz growth functions. Although there have been calls for moving beyond the von Bertalanffy and Gompertz growth functions for statistical (e.g., negative correlation between parameter estimates) and life-history (e.g., they do not account for the change in energy allocation after sexual maturity) reasons [32], they are still popular — and almost the default choices — among modelers.

#### 2.3.1 The von Bertalanffy growth function

The von Bertalanffy growth function (vBGF) has been used to model the growth of fish [33], mammals [34], snakes [35], birds [36] and many other species and taxa. von Bertalanffy hypothesized that the growth of an organism results from a dynamic balance between anabolic and catabolic processes [9]. If *W* (*t*) denotes mass at time *t*, the von Bertalanffy assumption is that anabolic factors are proportional to surface area, which scales as *W* (*t*)^2/3^, and that catabolic factors are proportional to mass. If *a* and *b* denote these scaling parameters, then the rate of change of mass is:

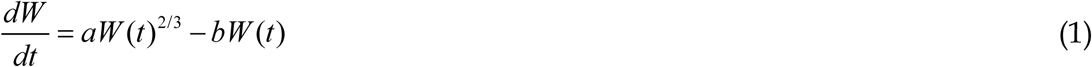

If we assume that mass and length (i.e., size), *L*(*t*), are related by *W* (*t*) = *ρ L*(*t*)^3^ with *ρ* corresponding to density, then calculus shows that:

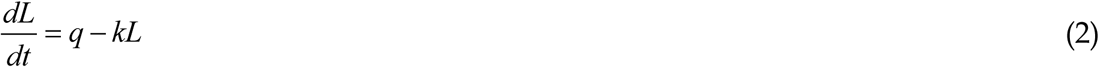

where *q* = *a / 3ρ* and *k* = *b / 3ρ*.

Setting 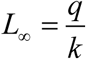 to be the asymptotic size and *L*(0) = *L*_0_ to be the initial size, two forms of the solution are:

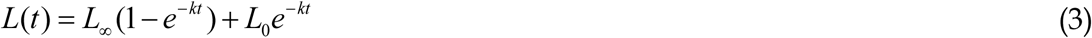

and:

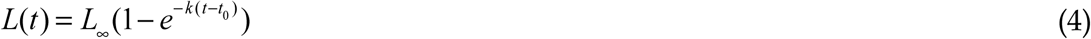

where *t*_0_ is the hypothetical age at which length is equal to 0.

If *L*(*t*) > *L*_∞_, the rate of growth *k* (in time^-1^ units) is negative, so asymptotic size is the upper limit of size, which is only attained in the limit of infinite time. In this work, we will use the formulation of the vBGF of Eq. 4, which has 3 parameters: *L*_∞_, *k*, *t*_0_. Although the definition of asymptotic size in the vBGF introduces an explicit linear relationship on the log scale between *k* and *L*_∞_ (i.e., log(*L*_∞_) = log(*q*) − log(*k*), according to Eq. 4), in this work we do not treat *L*_∞_ as equal to 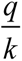 (see [37] for a formulation of the function explicitly including the link between *L*_∞_, *q*, and *k*), but we let the correlation between *L*_∞_ and *k* at the whole population and at the individual level emerge from data.

#### 2.3.2 The Gompertz growth function

The Gompertz growth function (GGF) [7, 8] has been used to model the growth of a variety of species and taxa, from plants to birds and fish growth, to tumor and bacterial growth [38, 39]. Contrary to the von Bertalanffy growth function, which was developed by von Bertalanffy from physiological principles, the Gompertz curve was first proposed by Benjamin Gompertz [8] for modeling mortality curves, and later adopted for studies of growth when it was empirically found that the GGF could well describe the growth trajectories of many species [7]. Therefore, the biological interpretation of the parameters of the Gompertz growth function is harder and more *a posteriori* than that of the von Bertalanffy function (but see [40] for an example of biological interpretation of the Gompertz model for tumor growth), and we introduce the parameters of the Gompertz growth function mostly as curve-fitting parameters.

The GGF has been used in a variety of parameterizations, with some formulations that have parameters that are more interpretable than in others [38]. One commonly found parameterizations for the GGF is:

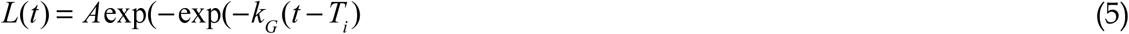

Where *L*(*t*) is size at time *t*, *A* (we might also call this parameter *L*_∞_ like in the von Bertalanffy growth function, since the two parameters both represent the asymptotic size), *k*_G_ (in time^-1^ units) is a coefficient of growth affecting the slope, and *T*_i_ is time at inflection, which in this formulation occurs when 37% of the final growth has been reached. *T*_i_ shifts the growth curve horizontally without changing it shape and therefore is a location parameter, while *A* and *k*_G_ are shape parameters.

### 2.4 Parameter estimation and individual variation

In this section, we will mostly make reference to the vBGF for describing mixed-effects models of growth and the estimation of their parameters, but all assumptions and methods are also applicable to the GGF and other similar size-at-age growth functions. In the vast majority of applications of the vBGF and of the GGF, *L*_∞_, *k*, and *t*_0_ (and *A*, *k*_G_, and *T*_i_) have been estimated at the population level (i.e., without accounting for individual heterogeneity in growth) starting from cross-sectional data, and interpreted as the growth parameters of an average individual in the population (i.e., *L*_∞_ is the asymptotic size of an average individual of the population or species). In fishery science, von Bertalanffy growth function’s *k* and *L*_∞_ and estimates of adult mortality are commonly used descriptors of the life-history strategies of fish populations [41]. Age-structured stock assessment methods are based on sizes-at-age estimations that are often derived from parameters of the von Bertalanffy growth model for that species [42].

[11] and [37] presented formulations of the vBGF specific for longitudinal data where L_∞_, *k*, and *t*_0_ may be allowed to be a function of shared predictors and individual random effects. Since *k* and *L*_∞_ must be non-negative, it is convenient to use a log-link function. We thus set for individual i in group j:

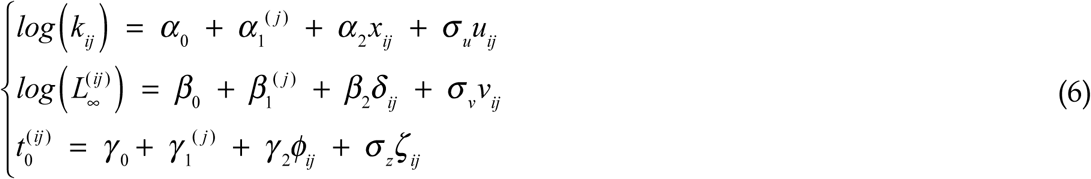

where ***u*** ∼ *N* (0,1), ***v*** ∼ *N* (0,1), and ***z*** ∼ *N* (0,1) are the standardized individual random effects, *σ_u_*, *σ_v_*, and *σ_z_* are the standard deviations of the statistical distributions of the random effects, 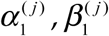, and 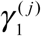 are group (i.e., categorical) effects (e.g., species, population, sex, year-of-birth), and the other parameters are defined as in Eq. 4. The continuous predictors *x_ij_*, *δ_ij_*, and *ζ_ij_* in Eq. 6 (e.g., population density, temperature, proxies of food quality and availability) do not need to enter linearly into the predictor, and the terms *β*_2_*δ_ij_*, *α*_2_ *x_ij_*, and*γ*_2_*ϕ_ij_* may be replaced by a more general function *f* (*x_ij_*; ***τ***), where ***τ*** denotes a set of parameters to be estimated. For the GGF, the parameters *A*, *k*_G_, and *T*_i_ are set the same way as in Eq. 5.

Length of individual *i* in group *j* at age *t* is for the von Bertalanffy function:

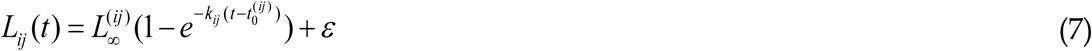

and for the Gompertz function:

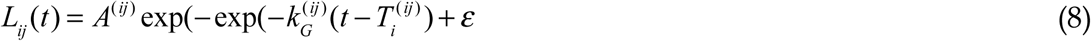

where *ε* is the observation error, which is time-invariant and normally distributed with mean 0 and variance 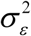.

According to Eqs. 7 and 8, a positive correlation between 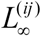 and *k_ij_* (from now on we will refer to them as *L*_∞_ and *k* at the individual level) and *A*^(*ij*)^ and 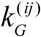 (*A* and *k_G_* at the individual level) indicates that size ranks tend to be maintained through the life of individuals, while a negative correlation indicates that size ranks tend not to be maintained [11, 37].

### 2.5 Model fitting

It has often been difficult, if not impossible using standard approaches, to estimate parameters for many of the proposed growth models using data on individual growth trajectories in natural settings (i.e., “in the wild”). In addition to noisy and sparse data, and even in the presence of a large amount of data, a highly parameterized non-linear model may be only weakly statistically identifiable. Several modeling tools to fit hierarchical models are now available, such as platform-independent BUGS [43], JAGS [44], Stan [45], and, among many others, the *nlme*, *lme4*, *NIMBLE*, *brms* packages and associated functions in R [19,46–48] or *PyMC3* in Python [49].

However, when dealing with a large number of random effects, non-linear models, and missing or noisy data, some of those methods may fail to converge or take a very long time to converge, thus limiting the number of explorations that can be carried out. Since many models are typically fitted when investigating biological process, it is always convenient, and often necessary in practice, to use algorithms and software that allow for rapid model exploration, for instance by reducing the time needed for the optimization algorithm to converge or by finding a compromise between obtaining the full posterior distribution of parameters (more computationally expensive) or only its summary statistics (less computationally expensive).

A tool that has been recently developed for fitting highly parameterized mixed-effects models, and which we used for fitting models in the present study, is Template Model Builder (TMB, [28]). TMB is a general random effect tool integrated in R that was inspired by ADMB (Automatic Differentiation Model Builder [50]), an open-source statistical software package for fitting highly parameterized non-linear statistical models with or without random effects. TMB can fit generic random-effects models using an Empirical Bayes approach that evaluates and maximizes the Laplace approximation of the marginal likelihood [51] using automatic differentiation. TMB computes standard errors of parameter estimates and of predictions using the delta method [52], offers easy access to parallel computations, and is very flexible in model formulation. Recent developments allow the estimation of parameters in Bayesian models using TMB [53].

### 2.6 Statistical analyses

Prediction of future observables has long been included as an aspect of statistics, but it has been much less prominent than parametric statistical inference [54]. More recently, it has been proposed that the proper use of statistical models is the prediction of future observations, and not uncertainty around estimates of model parameters [55]. In this work, we will evaluate models from the point of view of both inference (parameter estimates and their biological interpretations) and predictive performance.

Since exploratory investigations often prevent the use of null-hypothesis testing, and multiple comparisons increase the “researcher degrees of freedom”, including the choice of convenient hypotheses to test [56], apart from some specific tests of correlation, we present and discuss our results of model results from a qualitative point of view, that is without formal null-hypothesis testing.

Both the GFF and vBGF assume that size data are evenly spaced in time. For the analyses of growth, we thus used only September data (unique fish sampled in September: LIdri_MT, *n* = 784; UIdri_MT, *n* = 502, LIdri_RT, *n* = 109, UVol_BT, *n* = 2434). [21] found no or minor effects of population density, water temperature, or sex on growth in Slovenian populations of marble, rainbow, and brown trout. In this study, we pooled all data from different populations together and use only group variables as candidate factors for explaining variation in growth. We introduced group predictors as fixed effects (since treating a factor with just a few levels as “random” may generate imprecise estimates of the associated standard deviation [57]) to test whether they improved model performance with respect to a model with no predictors other than random effects. In particular, we included as predictors (i) *Species* (3-level predictor: marble trout – *MT*; brown trout – *BT*; rainbow trout – *RT*), and (ii) *Population (*4-level predictor: *LIdri_MT*, *UIdri_MT*, *LIdri_RT*, *UVol_BT*) as a group (i.e., categorical) variable. We tested for vBGF and GGF all the combination of *Species*, *Population*, and *Constant* (i.e., no predictors) for the three parameters of either growth function. We also experimented with *Cohort* (i.e., year-of-birth) as a group predictor and with two-way interactions between *Species*, *Population*, and *Cohort*.

We tested the predictive ability of the candidate vBGF and GGF models as follows. For each model, we first tested for convergence of the TMB algorithm and computed the Akaike Information Criterion (AIC [58]) when fitting the model on the whole data set. We then: (*i*) identified fish that were sampled more than 3 times; (*ii*) randomly sampled one third of them (test data set, 201 unique individuals); (*iii*) deleted from the data set all observations except the first one from each individual fish in the validation sample; (*iv*) if the TMB algorithm converged, estimated the parameters of the vBGF and GGF for each individual including those in the validation sample; (*v*) predicted the observations in the test data set (*n* between 522 and 541 from the 201 unique individuals); and (*vi*) estimated or computed accuracy measures, such as *R*^2^ with respect to the 1:1 prediction-observation line, and maximum error for the test sample. We repeated (*ii*)-(*vi*) 5 times and then computed the mean of the accuracy measures for each vBGF and GGF model.

Following [11] and [37], we estimated the correlation between *L*_∞_ and *k,* and *A* and *k_G_*, to get insights into potential processes driving the observed variation in growth. We also estimated the correlation between *L*_∞_ and *A* at the individual level within populations when using the same models for the two growth functions, since the values of *L*_∞_ and *A* are both estimates of asymptotic size.

## 3 Results

Results are fully reproducible. Data and R code are at https://github.com/simonevincenzi/Growth_Models.

There were 60 individuals sampled 3 or more times in LIdri_MT, 7 in LIdri_RT, 90 in UIdri_MT, and 513 in UVol_BT. Empirical growth trajectories showed substantial individual variation in growth rates and size-at-age (Fig. 1).

**Fig. 1.**
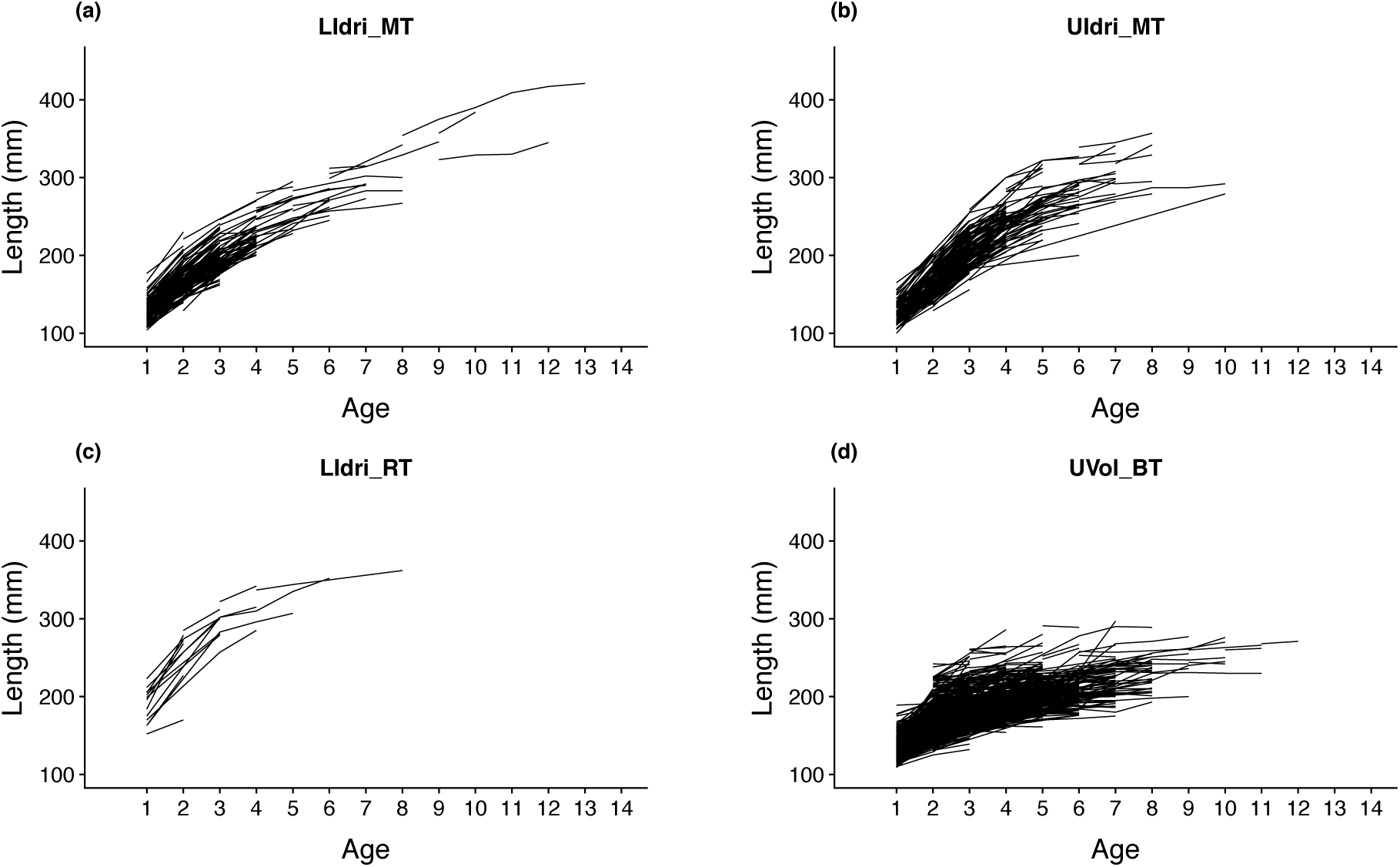
Growth trajectories of fish sampled more than once between 2004 and 2015 (only September samplings) in the populations of LIdri_MT (*n* = 210 unique fish), UIdri_MT (*n* = 209), LIdri_RT (*n* = 17), and UVol_BT (*n* = 1323).

Most models including either *Cohort* or interacting predictors never converged or converged only for some of the five replicates, and we thus dropped them from the set of candidate models. A number of different vBGF and GGF models had basically the same predictive accuracy, and their AICs computed on the full data set were fairly close as well (Table 1). For 12 out of the 24 models, *R*^2^ with respect to the 1:1 predicted-observed line was greater than 0.80 (Table 1). The overall correlation (Pearson’s *r*) between *L*_∞_ and *A* when using the same model (i.e., same predictors for the “equivalent” parameters) and pooling together all population-specific estimates was 0.51 (*p* < 0.01) (Fig. 2). However, within-population correlations between *L*_∞_ and *A* were all non-significant.

**Fig. 2.**
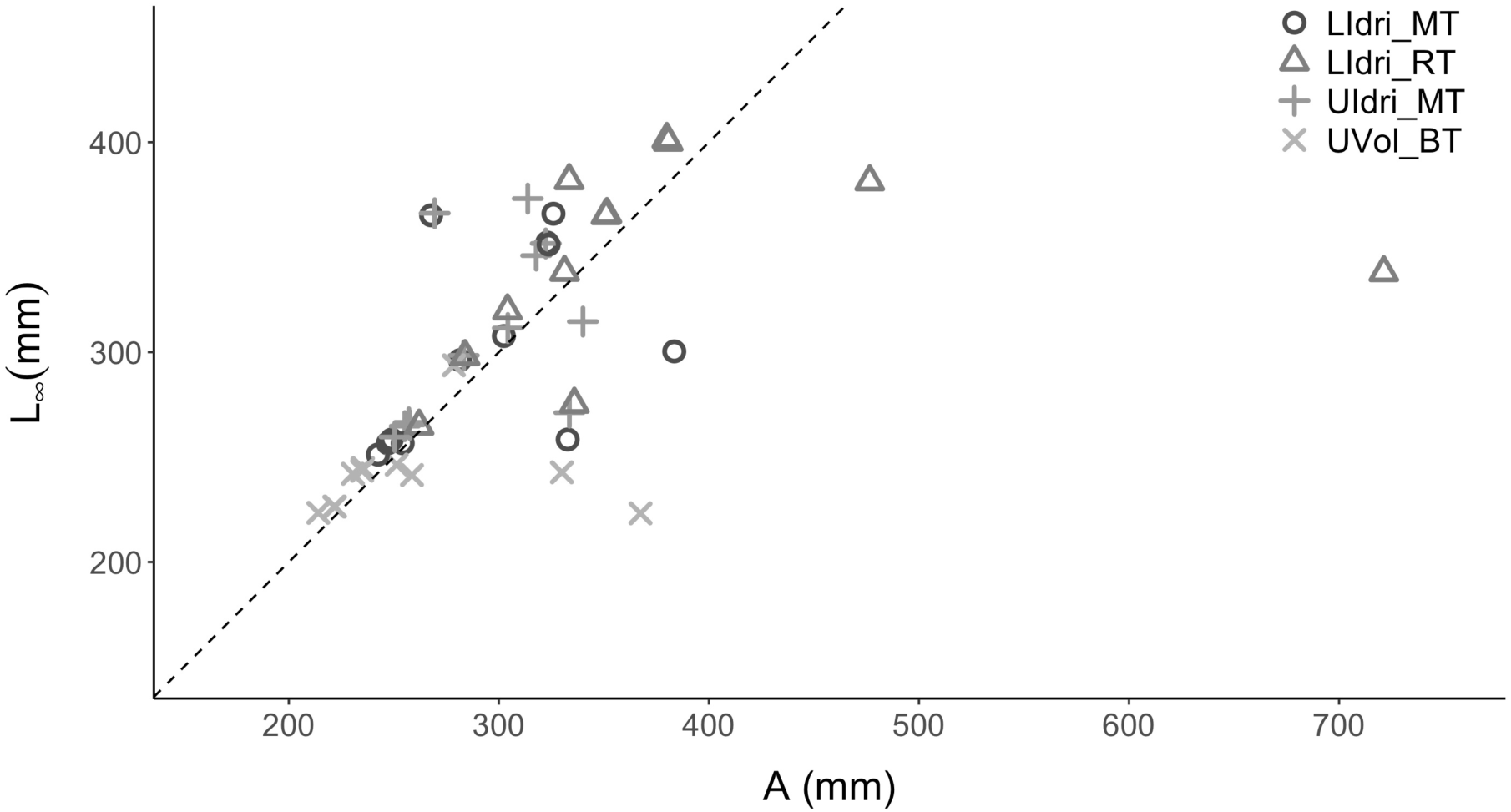
Estimates of *L*_∞_ and *A* for the same models (i.e., same predictors for the equivalent model parameters). The mean [min, max] of the ratio *L*_∞_*/ A* across the 12 models were 1.03 [0.78, 1.36] for LIdri_MT, 0.96 [0.47, 1.14] for LIdri_RT, 1.06 [0.13, 0.81] for UIdri_MT, and 0.96 [0.143, 0.60] for UVol_BT. The dashed line is the 1:1 line.

**Table 1.** Ranking of models according to their mean *R*^2^ with respect to the 1:1 predicted-observed line over five random test data sets. *R*^2^ (sd) is the standard deviation of the five *R*^2^ calculated for each model. In parentheses, the predictors for the growth function parameters, either *Species*, *Population* (*Pop*), or *Constant* (i.e., no predictors - *Const*). Function is either the von Bertalanffy (*vBGF*) or the Gompertz (*GGF*) growth function. The best model according to the Akaike Information Criterion (AIC) was the Gompertz growth function model with *A*(*Pop*), *k_G_* (*Pop*), *T*_i_ (*Pop*).

| **Model** | **Function** | **R<sup>2</sup> (mean)** | **R<sup>2</sup> (sd)** | **AIC** |
| --- | --- | --- | --- | --- |
| $L_{\infty}(\text{Species}), k(\text{Pop}), t_0(\text{Pop})$ | $vBGF$ | 0.852 | 0.02 | 54717.75 |
| $L_{\infty}(\text{Pop}), k(\text{Pop}), t_0(\text{Pop})$ | $vBGF$ | 0.851 | 0.02 | 54717.12 |
| $L_{\infty}(\text{Pop}), k(\text{Pop}), t_0(\text{Const})$ | $vBGF$ | 0.847 | 0.02 | 54771.37 |
| $A(\text{Species}), k_G(\text{Pop}), T_i(\text{Pop})$ | $GGF$ | 0.846 | 0.03 | 54749.06 |
| $L_{\infty}(\text{Species}), k(\text{Pop}), t_0(\text{Const})$ | $vBGF$ | 0.846 | 0.02 | 54775.51 |
| $A(\text{Pop}), k_G(\text{Pop}), T_i(\text{Pop})$ | $GGF$ | 0.846 | 0.03 | 54678.04 |
| $A(\text{Pop}), k_G(\text{Const}), T_i(\text{Pop})$ | $GGF$ | 0.845 | 0.02 | 56017.61 |
| $A(\text{Species}), k_G(\text{Const}), T_i(\text{Pop})$ | $GGF$ | 0.843 | 0.02 | 54768.57 |
| $L_{\infty}(\text{Pop}), k(\text{Const}), t_0(\text{Pop})$ | $vBGF$ | 0.842 | 0.02 | 54942.05 |
| $L_{\infty}(\text{Species}), k(\text{Const}), t_0(\text{Pop})$ | $vBGF$ | 0.838 | 0.02 | 54962.76 |
| $L_{\infty}(\text{Const}), k(\text{Pop}), t_0(\text{Pop})$ | $vBGF$ | 0.810 | 0.02 | 55546.41 |
| $A(\text{Const}), k_G(\text{Pop}), T_i(\text{Pop})$ | $GGF$ | 0.801 | 0.02 | 55647.87 |
| $A(\text{Pop}), k_G(\text{Pop}), T_i(\text{Const})$ | $GGF$ | 0.770 | 0.03 | 56047.32 |
| $A(\text{Species}), k_G(\text{Pop}), T_i(\text{Const})$ | $GGF$ | 0.768 | 0.03 | 56222.17 |
| $L_{\infty}(\text{Pop}), k(\text{Const}), T_i(\text{Const})$ | $vBGF$ | 0.714 | 0.04 | 56551.44 |
| $A(\text{Pop}), k_G(\text{Const}), T_i(\text{Const})$ | $GGF$ | 0.712 | 0.04 | 56568.42 |
| $L_{\infty}(\text{Species}), k(\text{Const}), t_0(\text{Const})$ | $vBGF$ | 0.708 | 0.04 | 56577.43 |
| $A(\text{Species}), k_G(\text{Const}), T_i(\text{Const})$ | $GGF$ | 0.707 | 0.04 | 56596.06 |
| $L_{\infty}(\text{Const}), k(\text{Const}), T_i(\text{Pop})$ | $vBGF$ | 0.700 | 0.06 | 56767.01 |
| $A(\text{Const}), k_G(\text{Pop}), T_i(\text{Const})$ | $GGF$ | 0.700 | 0.05 | 56843.89 |
| $A(\text{Const}), k_G(\text{Const}), T_i(\text{Pop})$ | $GGF$ | 0.698 | 0.06 | 57097.36 |
| $L_{\infty}(\text{Const}), k(\text{Const}), t_0(\text{Const})$ | $vBGF$ | 0.679 | 0.05 | 57567.55 |
| $A(\text{Const}), k_G(\text{Const}), T_i(\text{Const})$ | $GGF$ | 0.675 | 0.05 | 57570.43 |
| $L_{\infty}(\text{Const}), k(\text{Pop}), t_0(\text{Const})$ | $vBGF$ | 0.665 | 0.06 | 56896.74 |

The GGF model with *Population* as predictor for all 3 parameters was the model with the best AIC computed on the whole data set (ΔAIC with the second-best model = 39.08) (Table 1). The equivalent-in-predictors vBGF model showed estimates of asymptotic size that were similar only for UVol_BT to those provided by the Gompertz growth function (Fig. 3). In both vBGF and GGF models with *Population* as predictor for all 3 parameters, *L*_∞_ and *k*, and *A* and *k_G_* at the individual level were positively correlated within populations (Fig. 4).

**Fig. 3.**
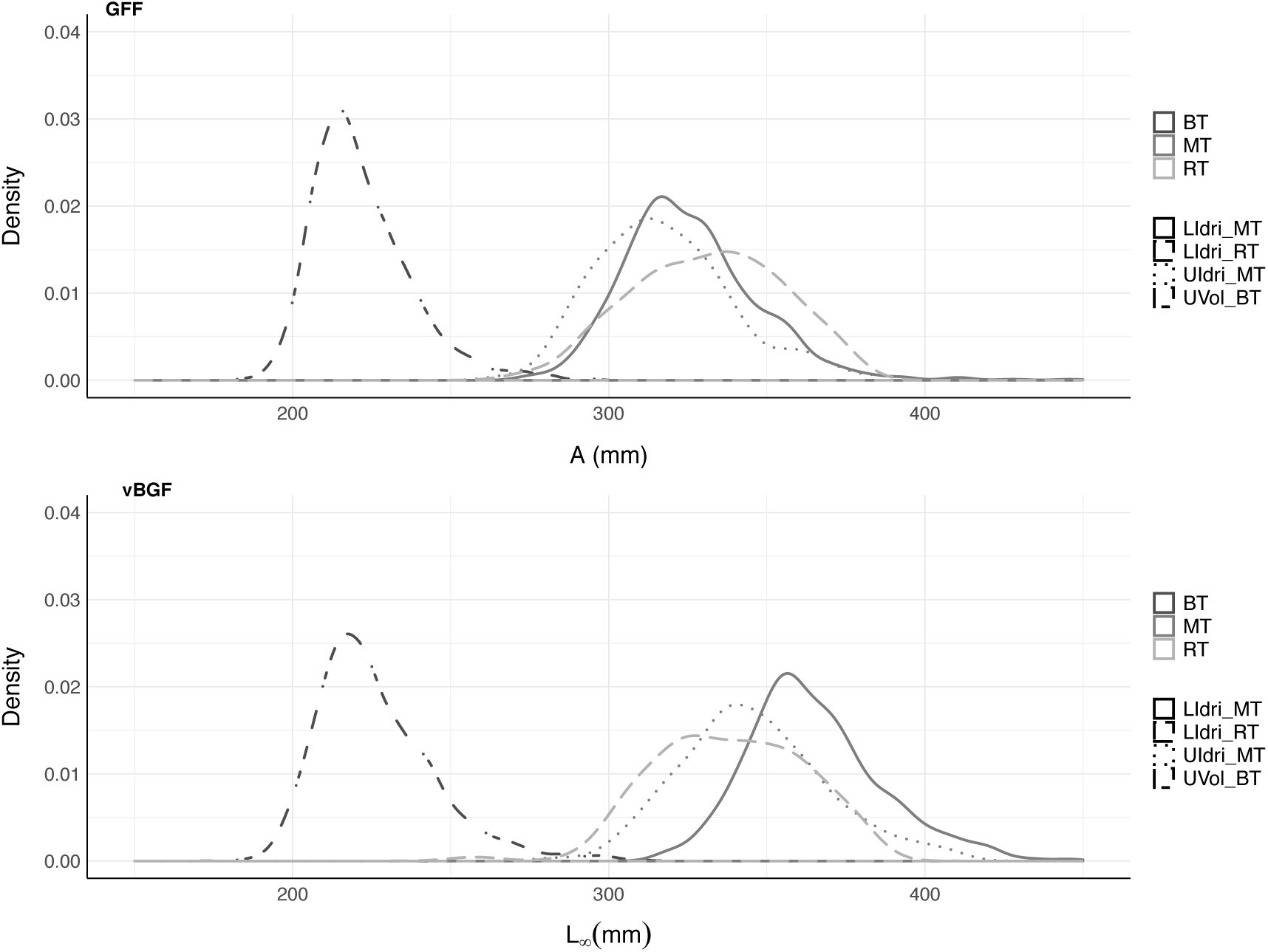
Empirical distribution of population-specific individual *L*_∞_ and *A* for the best Gompertz growth function *GGF* (and equivalent von Bertalanffy growth function *vBFG*) model according to AIC (i.e., *Population* predicting all 3 parameters). *GGF* (Mean (sd)): LIdri_MT, 326.00 mm (22.52); UIdri_MT, 317.79 (22.71); LIdri_RT, 331.29 (23.24), UVol_BT, 221.66 (15.61). *vBGF*: LIdri_MT, 365.93 mm (23.82); UIdri_MT, 346.00 (24.11); LIdri_RT, 338.15 (23.22), UVol_BT, 226.44 (18.70).

**Fig. 4.**
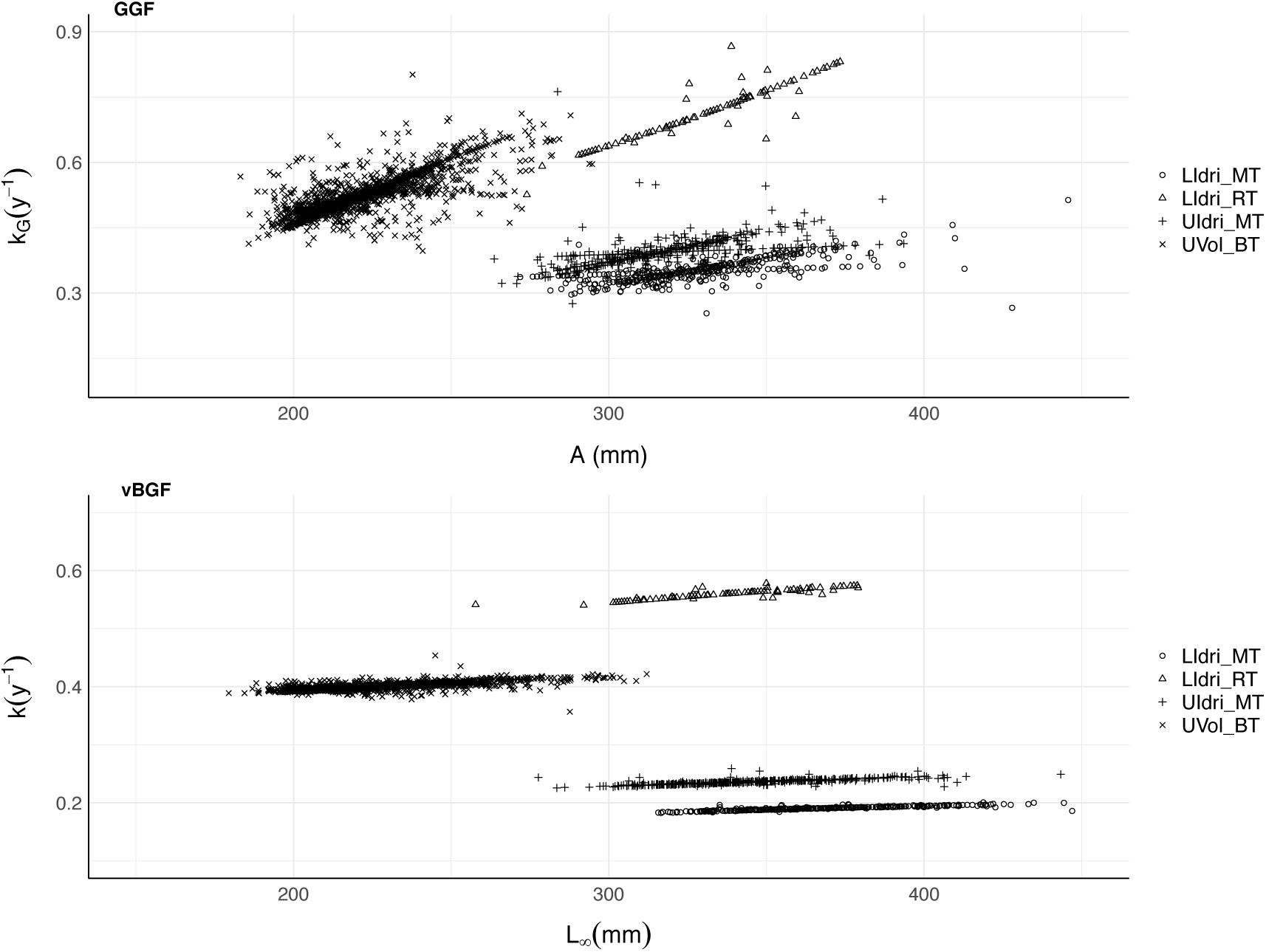
Correlation between *L*_∞_ and *k* for the *vBGF* and *A* and *k_G_* for the *GGF* for the best Gompertz growth function *GGF* (and equivalent von Bertalanffy growth function *vBFG*) model according to AIC (i.e., *Population* predicting all 3 parameters). *GGF* (Pearson’s *r, all p < 0.01)*: LIdri_MT (*r* = 0.65); UIdri_MT (0.68); UVol_BT (0.63); LIdri_RT (0.33). *vBGF*: LIdri_MT (*r* = 0.90); UIdri_MT (0.83); UVol_BT (0.71); LIdri_RT (0.95).

GGF and vBGF’s predictions of sizes in the test data sets were remarkably similar to each other, with Pearson’s correlation between GGF and vBGF predictions for the same models (i.e., same predictors used in either growth function) on average greater than 0.99 (both prediction and parameter estimate uncertainties are provided in the online resources associated with this paper).

However, despite the strong positive correlation between GGF and vBGF model predictions, for the model with *Species* predicting *A* and *L*_∞_ for GGF and vBGF, and *Population* predicting the other two parameters for either function (i.e., the best model in term of average predictive accuracy on test data sets for both the GGF and vBGF, see Table 1), the estimated A and *L*_∞_ were largely different from each other for the population of LIdri_RT 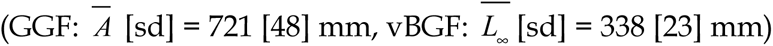 and similar for the other 3 populations (Fig. 5). In addition, for LIdri_RT the correlation between *A* and *k_G_* in the GGF was negative (*r* = −0.56, *p* < 0.01), and between *L*_∞_ and *k* in the vBGF was positive (*r* = 0.91, *p* < 0.01).

**Fig. 5.**
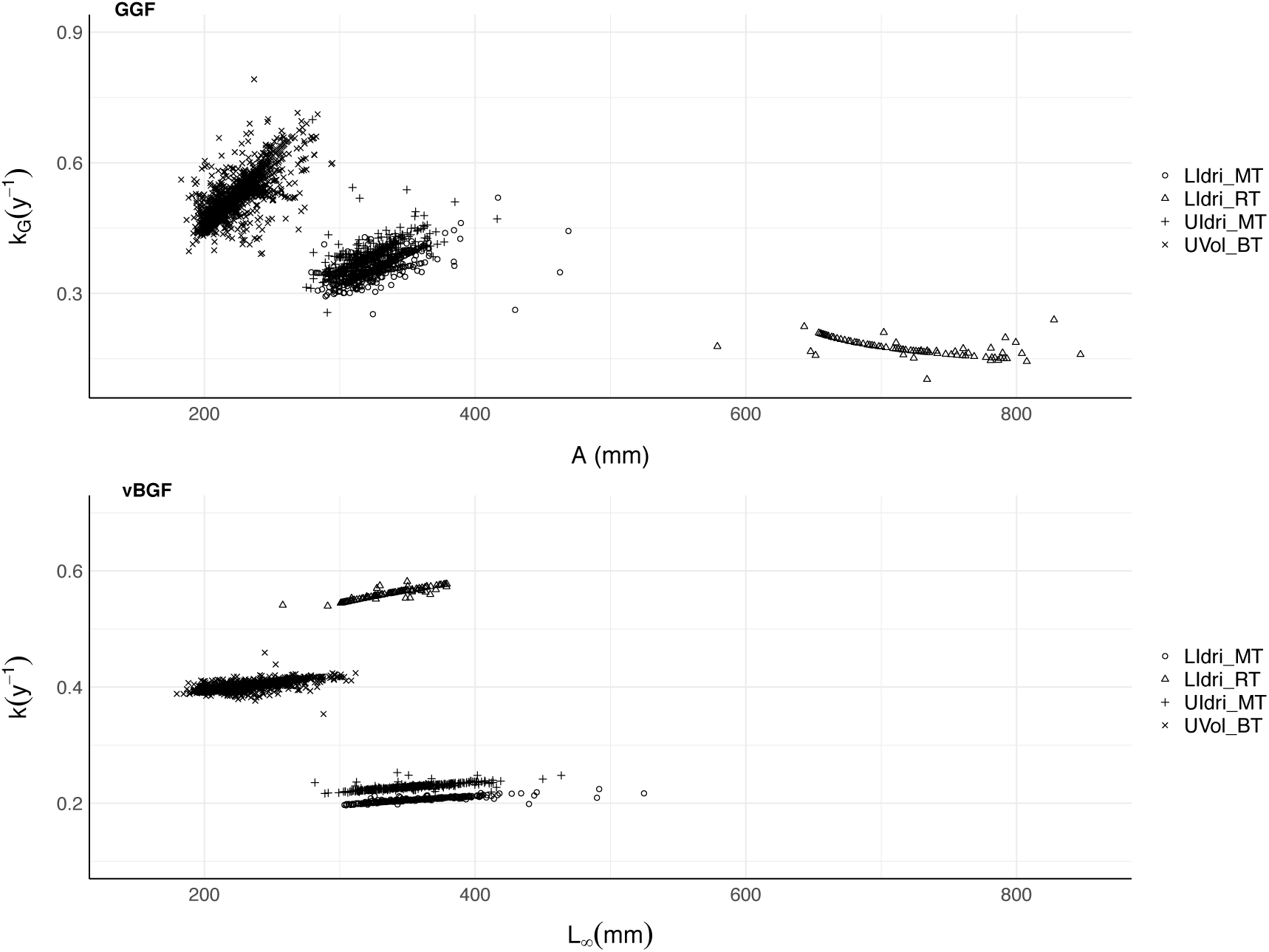
Correlation between *L*_∞_ and *k* for the *vBGF* and *A* and *k_G_* for the *GGF* for the best von Bertalanffy growth function *vBFG* (and equivalent Gompertz growth function *GGF*) model according to prediction performance (i.e., *Species* predicting asymptotic size, and *Population* predicting the other 2 parameters). *GGF* (Pearson’s *r, all p < 0.01)*: LIdri_MT (*r* = 0.65); UIdri_MT (0.59); UVol_BT (0.76); LIdri_RT (−0.56). *vBGF*: LIdri_MT (*r* = 0.91); UIdri_MT (0.81); UVol_BT (0.73); LIdri_RT (0.91).

Across models, a few individuals living in UVol_BT and UIdri_MT were consistently those with the largest prediction errors. Those individuals had growth trajectories that were unusual with respect the common growth trajectories in their populations (Fig. 6). Both in the many trajectories that were predicted accurately and in the few that were not, the growth trajectories estimated by the GGF and vBGF models with the same predictors were basically indistinguishable (Fig. 7).

**Fig. 6.**
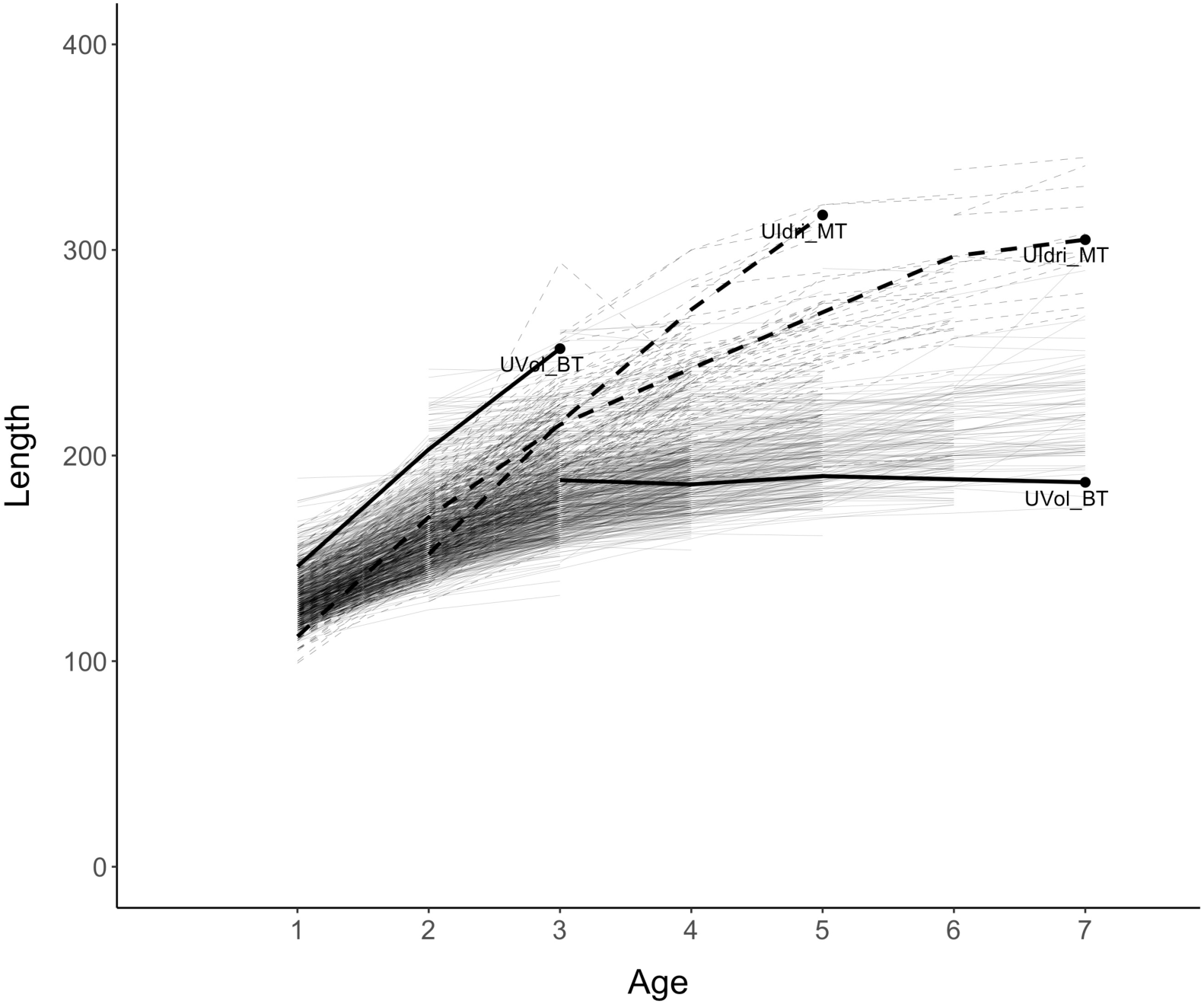
Growth trajectories of the 4 individuals (2 from UIdri_MT and 2 from UVol_BT) that were most consistently the worst predicted (the black circle is the worst prediction for the individual) by both vBGF and GGF models. Light dashed lines are the empirical trajectories of all fish sampled in UIdri_MT and light solid lines are the empirical trajectories of all fish sampled in UVol_BT.

**Fig. 7.**
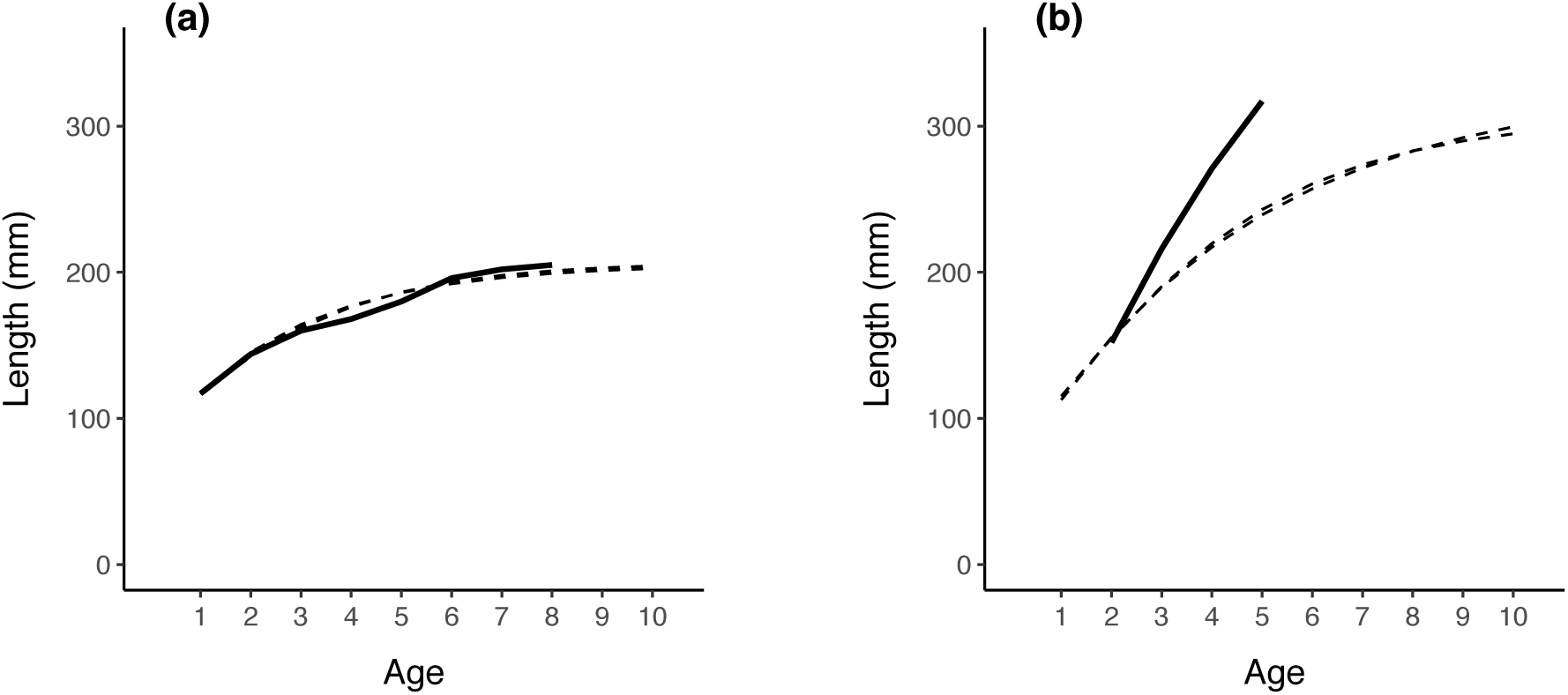
Example of good (a) and bad (b) predictions of growth (dashed lines) for GGF and vBGF models (all three parameters in either growth function with *Population* as predictor) for two individuals that have been sampled multiple times in UVol_BT (solid lines). The predictions of size-at-age provided by the GGF and vBGF were basically equivalent and based on size measured only at first sampling, at age 1+ for the individual in panel (a) and at age 2+ for the individual in panel (b).

## 4 Discussion

We found that mixed-effects models based on either the von Bertalanffy and Gompertz growth functions were able to largely capture the individual variation in growth among fish living in four distinct freshwater salmonid populations. Among the models we tested, the predictive performances on test data sets of the best von Bertalanffy and Gompertz mixed-effects models were largely equivalent, although their estimates of asymptotic size were often substantially different. Finally, parameter estimates for both growth functions point to strong maintenance of size hierarchies over time in each of the four salmonid populations.

### 4.1 Growth processes

Individuals living in the same environment, and especially those of species with growth after sexual maturity, often vary in their body growth rate and size-at-age. In the four trout populations that have been investigated in this work, the size of the smallest age-1 fish was ∼50% of the size of the biggest age-1 fish. Growth trajectories in fish are often consistent through time, that is individuals that are small early in life are likely to be among the smallest years later [11, 59]. [11] showed that a positive correlation between asymptotic size and growth rate points to the maintenance of size hierarchies through the lifetime of organisms, that is if individual *a* is bigger than individual *b* at age *t*, individual *a* is likely to be bigger than individual *b* at time *t*+1…*t*+n. For all four populations, we found a positive correlation between asymptotic size and growth rate at the individual level when using either growth function.

In freshwater trout, the primary type of intra-specific competition for resources seems to be interference competition for space [37], probably due to their strong territoriality. In interference competition, bigger individuals reduce the access to resources, such as space and food, of smaller individuals, and may also live longer. High heritability of growth [25], maternal decisions on the timing and location of spawning [60], and dominance established early in life [61] are all processes that in combination or by themselves may explain the maintenance of size ranks throughout fish lifetime.

### 4.2 Prediction of growth trajectories

There is a rich literature on the comparison between growth function for prediction of unobserved data and inference on growth processes in species [62–65]. When developing mathematical and statistical models in biology and ecology, in particular when those models are used for making predictions of unobserved or future realizations of biological processes, we face trade-offs between model complexity, interpretability of model parameters, ease of parameter estimation, and accuracy of predictions. It is common for more complex models either in number of predictors, how the predictors enter the model (e.g., non-linearly) or the algorithm used to estimate model parameters, to provide higher accuracy (here, both the ability of the model to explain observed data and make correct predictions about future observables). However, higher accuracy may come at the expense of ease of parameter estimation, interpretability of model predictors and parameters (i.e., to what degree the model allows for understanding processes), and costs of collecting predictor values or maintaining data pipelines.

The variation in growth and size that characterizes organisms can almost always be modeled retrospectively, but size-at-age is more difficult to forecast. The limited number of attempts at predicting missing size observations or unobserved size-at-age and growth trajectories may also depend on the intrinsic unpredictability of some growth curves. For instance, in species with environmental sexual determination and sexual dimorphism, such as eels [66], or when growth is faster later in life and is strongly determined by the environment (e.g., ocean growth of anadromous salmonids [67]), it may be impossible to accurately predict later portions of the growth trajectory when only observations early in life are available.

We have shown that for the four salmonid populations that we used as a model system, the best Gompertz and von Bertalanffy mixed-effects growth models allow one to use a single measurement early in the life of individual fish to obtain accurate predictions of their size-at-age in the future. In addition, the predictions made by the two models when using the same predictors for their parameters were basically the same; choosing between the best models of the two growth functions would have very few practical consequences for predictions of size-at-age of individuals. Some inaccurate predictions deserve further investigations, although they are to be expected when predicting the realizations of complex biological processes.

However, estimates of asymptotic size within populations were not correlated when estimated using the same Gompertz and von Bertalanffy growth models. For instance, for one highly ranked model in terms of AIC and predictive performance for both growth functions, the estimates of asymptotic size for rainbow trout provided by the Gompertz model were on average more than two times bigger than those provided by the von Bertalanffy model; in addition, for the same model, asymptotic size and growth rate at the individual level for rainbow trout were negatively correlated when using the Gompertz model and positively correlated when using the von Bertalanffy model.

Berkey (1982) found that growth curve parameters estimated using the Empirical Bayes method (although the same can be said of any estimation method for size-at-age models) are particularly sensitive to the end points of the growth trajectories. In our study, the population of rainbow trout had the smaller and sparser data among all four salmonid populations we used as a model system. However, wildly different combinations of parameters of these growth functions can result in some cases in very similar growth trajectories over a restricted time horizon [11]; this explains why the resulting predictions provided by the two models were basically the same, although the inference on growth processes coming from the analysis of parameter estimates was different.

Although predictions from models of the two growth functions with similar performance on test data sets were highly correlated and barely distinguishable, the two growth functions typically provided different (in some cases, substantially different) estimates of asymptotic size. This suggests that distribution of size-at-age can be a more informative and stable-across-models measure and descriptor of growth than estimates of model parameters, especially when using or comparing different growth functions.

Shohoji et al. (1991) found that the classification of individuals into homogenous groups (i.e., where the strength is borrowed from in mixed-effects models) was necessary to obtain accurate predictions of human lifetime growth. In our case, the classification either into population or species was sufficient to develop models that provided overall excellent predictions of the future growth of fish. When a single measurement early in life is sufficient to make accurate predictions of future growth, we can hypothesize that either the intrinsic growth potential of the individual, the environment experienced early in life, or a combination of the two largely determine the lifetime growth of individuals. Strong empirical evidence of early induced effects on later-in-life growth rate, life-history traits, and behavior of organisms is rapidly building up in the literature [70, 71].

## 5 Conclusion

A better understanding of the evolutionary, physiological, and life-history determinants of the differences in growth within and between individuals, populations, species, and taxa will always a major problem in biology. In the context of species that grow after sexual maturity and when used for predictive purposes, it appears that size-at-age mixed-effects growth models with time-invariant predictors sit in a favorable place on the surface that trades off accuracy, complexity, and biological interpretation of model parameters.

## Acknowledgements

We thank the MAVA Foundation for financial support and Marc Mangel for comments on the manuscript.

## Data Statement

### Data and R code

The file Readme.md in the repository https://github.com/simonevincenzi/Growth_Models describes each step of the reproducible analyses.

